# Hap10: reconstructing accurate and long polyploid haplotypes using linked reads

**DOI:** 10.1101/2020.01.08.899013

**Authors:** Sina Majidian, Mohammad Hossein Kahaei, Dick de Ridder

## Abstract

**Background:** Haplotype information is essential for many genetic and genomic analyses, including genotype-phenotype associations in human, animals and plants. Haplotype assembly is a method for reconstructing haplotypes from DNA sequencing reads. By the advent of new sequencing technologies, new algorithms are needed to ensure long and accurate haplotypes. While a few linked-read haplotype assembly algorithms are available for diploid genomes, there are no algorithms yet for polyploids.

**Results:** The first haplotyping algorithm designed for 10X linked reads generated from a polyploid genome is presented, built on a typical short-read haplotyping method, SDhaP. Using the input aligned reads and called variants, the haplotype-relevant information is extracted. Next, reads with the same barcodes are combined to produce molecule-specific fragments. Then, these fragments are clustered into strongly connected components which are then used as input of a haplotype assembly core in order to estimate accurate and long haplotypes.

**Conclusions:** Hap10 is a novel algorithm for haplotype assembly of polyploid genomes using linked reads. The performance of the algorithms is evaluated in a number of simulation scenarios and its applicability is demonstrated on a real dataset of sweet potato.

## 1. Background

Polyploids are organisms that possess three or more copies of each chromosome. There are numerous cases of polyploidy in the animal kingdom, including fish, amphibians and reptiles [1]. In plants, economically important crops such as potato, wheat, cotton and oat are polyploids [2]. For many genetic and genomic analyses, it is essential to know the sequence of alleles at variant sites corresponding to each homologous chromosome, i.e. the haplotypes. Haplotype information is needed to understand recombination patterns and uncover genotype-phenotype associations, with important applications in medicine [3] and plant breeding [2]. The development of DNA sequencing technologies, specific protocols and computational tools make it possible to reconstruct the haplotypes of individuals to some extent. Nevertheless, obtaining haplotypes of polyploids remains a challenging computational problem [4].

Several algorithms for polyploid haplotyping have been developed in recent years. In the absence of DNA sequencing errors, the haplotyping problem reduces to a simple clustering. If errors have to be taken into account, no polynomial-time solution exists. Therefore, different approximative and heuristic approaches have been used to estimate haplotypes. HapTree [5] is a greedy likelihood-based algorithm in which SNPs are added incrementally while keeping the tree of possible solutions to a manageable size. SDhaP [6] solves a correlation clustering problem using a gradient method to estimate the haplotypes. H-PoP [7], a heuristic algorithm, solves a combinatorial optimization problem called “polyploid balanced optimal partition”. Another approach is to use the minimum fragment removal (MFR) model in which conflicting fragments (due to erroneous reads) are removed. Siragusa *et al*. devised a new algorithm based on the MFR model, which uses integer linear programming [8].

The above-mentioned algorithms are developed solely for short reads generated by Illumina DNA sequencing machines. These produce reads that have a low sequencing error rate (~0.1%) but do not provide long-range information, which is key in reconstruction of long haplotypes. Over the last years, a novel category of sequencing technology characterized by long-read sequencing was developed and commercialized by Pacific Biosciences and Oxford Nanopore [9]. However, successful application of long-read sequencing for haplotyping is hampered by the still high sequencing error rate and significant costs involved.

Recently, 10X Genomics developed a linked-read sequencing library preparation strategy, commercialized through their Chromium platform, as a complementary technology to Illumina devices. This platform has the potential to provide long fragments at both low error rate and cost. In brief, the input genomic DNA, as little as 1 ng, is sheared into molecules of ~10–100 kbp. Subsequently, these molecules are isolated, partitioned into fragments, tagged with a unique 16bp barcode, and amplified on beads in an emulsion. The resulting material is then sequenced by normal Illumina paired-end technology, which results in high-throughput reads that contain long-range genomic information through these barcodes [10]. Such linked reads make it possible to assemble repetitive genomic regions as well as reconstruct long haplotype blocks. 10X Genomics delivers a likelihood-based algorithm to reconstruct haplotypes of diploid organisms such as humans in a software package called LongRanger [10–11]. However, no polyploid haplotyping algorithm is available at this moment, precluding the application of 10X-based haplotyping to a number of commercial crops and animals. Current polyploid haplotyping algorithms can be used on the obtained reads, ignoring the barcode information, but obviously the reconstructed haplotype blocks would be shorter than possible.

Exploiting the barcode information for haplotyping is possible by leveraging the so-called “fragment file” format. This format is used in preprocessing steps in several haplotyping algorithms [6, 12–13]. The extractHAIRs (Extract HAplotype Informative Reads) program in the HapCUT2 package [12] can be used to produce a fragment file based on aligned reads and heterozygous SNPs. While extractHAIRs is dedicated to diploids and is used for haplotyping based on 10X linked reads, the same concept (with some modifications) may be applied to polyploids. Using the obtained fragment file as input of a haplotype assembly core, SDhaP [6], long haplotype blocks of a polyploid can be reconstructed. However, in our simulations for a small genome using the aforementioned approach we obtained poor results in terms of reconstruction rate and vector error rate. Moreover, SDhaP crashes for larger datasets. This indicates that this short-read haplotyping algorithm is currently unable to directly handle 10X data generated from a polyploid genome.

To tackle this computational problem, we designed Hap10 – a first haplotyping software package specifically tailored for 10X linked reads generated from a polyploid genome. We provide a general framework based on SDhaP that allows haplotyping at the chromosome scale. Furthermore, we propose a novel optimization method that generates more accurate haplotypes with almost the same block length.

## 2. Methods

We have developed the Hap10 package to reconstruct haplotypes from a polyploid genome using 10X linked reads. Prior to haplotyping, several processing steps on sequencing reads are required. These include barcode handling, read alignment and variant calling, which are discussed in Section 2.1. Thereafter, Hap++, a new pipeline for polyploid haplotyping of 10X linked reads is explained in detail in Section 2.2. This pipeline uses SDhaP as the assembly core. Lastly, the Hap10 algorithm is presented in Section 2.3. This algorithm leverages the Hap++ pipeline, supplemented with a novel optimization based on an augmented Langrangian formulation as the assembly core. Section 2.4 concludes by discussing the data and performance measures used for validation of the method.

### 2.1 Preparation procedure

First, the 16bp 10X barcode is removed from the beginning of each paired-end read generated by the Illumina device. The barcode is stored as a read tag for further use. The possibility of sequencing errors in the barcode calls for an error correction scheme based on the known set of barcodes. Next, the reads are aligned to the reference genome using the barcode information, which provides a better alignment, particularly in repetitive genomic regions. These steps are performed using the LongRanger package (version 2.2.2) [11] provided by 10X Genomics, which generates a binary sequence alignment (BAM) file in which the barcodes are stored in the BX tag of each read. Subsequently, single nucleotide polymorphism (SNP) sites and their genotypes are called using the FreeBayes package (version 1.3.1) [14] with “-p 3” and “-p 4” for triploids and tetraploids, respectively and stored as a variant call format (VCF) file. The pipeline is depicted in Fig. 1.

**Figure 1.**
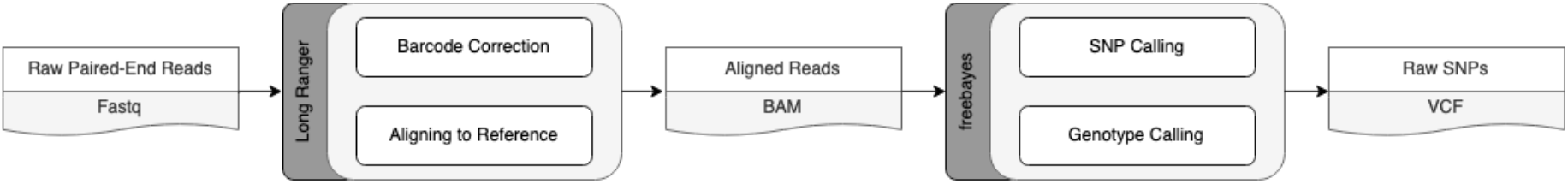
Preparation procedure for haplotyping of linked read data: barcode correction, read alignment and SNP/genotype calling. The output consists of aligned reads (BAM file) and called variants (VCF file).

### 2.2 Hap++

Hap++ is a fast program to reconstruct haplotypes in polyploids by exploiting linked read information. It consists of three main steps:

1. extracting haplotype-relevant information from input BAM and VCF files;
2. extracting molecule-specific fragments;
3. extracting strongly connected components of fragments.

The output of the last step can then be used by SDhaP to assemble the haplotypes. The three steps are described below.

#### Step 1. Extracting haplotype information

We first extract data relevant for haplotyping from the BAM and VCF files. As only heterozygous SNPs are informative for haplotyping, we filter out the homozygous variants from the VCF file. Next, we remove reads that cover fewer than two SNPs, since these do not provide any information for haplotyping. Subsequently, we extract the alleles of SNP sites of each read stored in the BAM file. In order to exploit long-range information provided by the barcodes, we combine the obtained fragments originating from the same 10X bead, i.e. with the same barcode. This results in long barcode-specific fragments. If there are two mismatching alleles for a SNP site corresponding to a specific barcode, we choose the one with the higher base quality. The result is a compact fragment file, similar to the output file of extractHAIRS [12, 15].

#### Step 2. Extracting molecule-specific fragments

The reads generated from the molecules in the 10X bead have identical barcodes. In an ideal case, the microfluidic device is expected to produce one molecule within each bead. In reality however there are, on average, 10 molecules per bead that originate randomly from one of the haploid chromosomes [10]. Therefore, the haplotypic origin of molecules with the same barcode is not identical, as discussed in [12]. As a result, parts of fragments in the fragment file are derived from different haplotypes, which misleads the haplotype assembly program. To tackle this issue, we propose a fragment processing scheme to extract molecule-specific fragments from each barcode-specific fragment. To this end, we use the mean-shift clustering algorithm [16] by means of its Python implementation from the Scikit-learn package [17]. We set the bandwidth of clustering to half of the expected 10X molecule length. This approach is based on the fact that molecule coverage is very low, and thus, molecules with the same barcode are generally distant from each other.

#### Step 3. Extracting strongly connected components of fragments

It is crucial to have a decent reference genome, because read alignment to the reference is upstream of haplotyping (Section 2.1). However, in practice, reference genomes are incomplete and contain gaps (usually represented by Ns). This affects haplotyping: if the reference contains a gap with length comparable with that of the 10X molecules, only few fragments connect the two sides of the gap and sequencing/mapping errors can have undue influence on the haplotyping process.

To prevent such problems, we first create a graph *G* in which fragments are considered as vertices *v* ∈ *G*. The weight *w*_*ij*_ of the edge *e*_*ij*_ = (*v*_*i*_, *v*_*j*_) between two nodes is calculated as the number of shared SNPs between two corresponding fragments, inspired by SDhaP [6]. As a demonstration, we generated such a graph for a read dataset (depicted in Supplementary information: Fig. S1), in which the length of the 10X DNA molecules is slightly higher than 50 kb, the length of the simulated gap. In this graph, one edge was found to connect two separate parts. This is based on a single molecule covering two distant SNPs in the vicinity of the gap. However, a single barcoded fragment is not enough for linking all haplotypes. Consequently, the accuracy of the whole haplotype block decreases.

As errors other than those due to gaps can lead to spurious edges in *G*, we provide a generic solution based on extracting strongly connected components of fragments. To this end, we exploit an iterative bipartitioning method based on the normalized cut (*NC*) [18]. We calculate the normalized Laplacian matrix (*L*_*N*_) of *G* based on the corresponding weight matrix *W*:

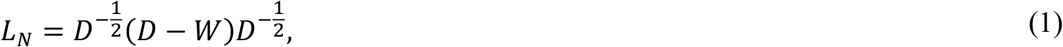

where *D* is the degree matrix of the graph, a diagonal matrix with *D*_*ii*_ = ∑_*j*_*w*_*ij*_. After calculating the eigenvalue decomposition of *L*_*N*_, we use the eigenvector (*E*_2_) that corresponds to the second smallest eigenvalue in order to bipartition the graph. The sign of the elements of vector *E*_2_ indicates the affiliation of fragments to either subgraph *G*_1_ or *G*_2_. Then, we calculate the *NC* value:

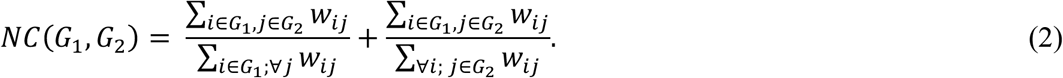

If *NC* is greater than a pre-specified threshold *t*, we stop the bi-partitioning procedure; otherwise, we continue bi-partitioning for each remaining partition. We set *t* to 0.03 for all simulations throughout the paper. When this step is finished, we output all strongly connected components of fragments as individual fragment files for processing by the assembly core, SDhaP. The Hap++ pipeline is depicted in Fig. 2. Note that this pipeline can be parallelized; specifically, the assembly core can be run on each strongly connected fragment simultaneously.

**Figure 2.**
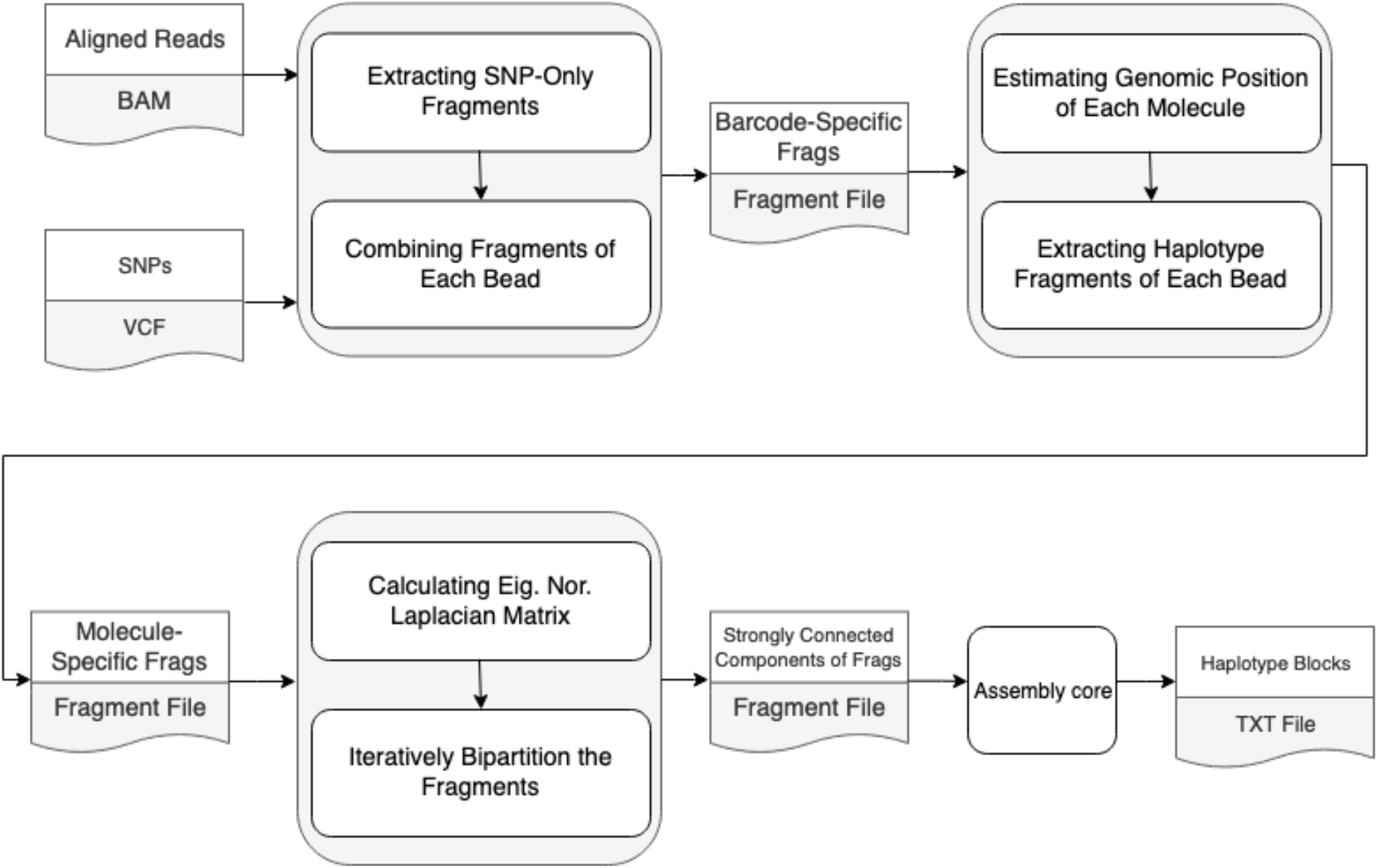
Hap++ pipeline. The output of the preparation procedure – BAM and VCF files – is pre-processed to make the haplotyping of 10X data feasible for polyploids. Next, strongly connected components of the molecule-specific fragment graph are extracted and used as input to the assembly core, which yields the haplotype blocks.

### 2.3 Hap10: an improved assembly core

The Hap10 pipeline leverages the Hap++ pipeline and adds a novel optimization as the assembly core. The goal of a haplotype assembly algorithm is to reconstruct *K* haplotypes *H* = {*h*_1_, … , *h*_*K*_} from *N* aligned fragments *R* = {*r*_1_, … , *r*_*N*_} generated by DNA sequencing of a *K*-ploid organism. This definition is universal and applies to different sequencing data types. Each *r*_*i*_ is assumed to originate from a single haplotype, as is the case for Illumina reads. As we discussed earlier, in 10X technology, we use molecule-specific fragments as *r*_*i*_ in our pipeline.

As a basis for Hap10, we use the three-step approach introduced by SDhaP:

I. Construct a fragment graph (similar to that of Section 2.2, Step 2) with weights between fragments (vertices) *i* and *j* calculated as.

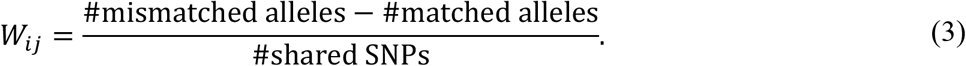
II. Split the fragments into *K* clusters, exploiting the graph weights.
III. Combine fragments of each cluster into a single haplotype using majority voting.

The reconstructed haplotypes are reported in a text file in a format similar to HapCUT2’s output presented in Supplementary information: Table S1.

Here, we explore step II of the assembly core. We use max-*K*-cut modelling for clustering the graph based on the edge weights *W*, which results in the following convex optimization problem osver *X* ∈ ℝ^*N*×*N*^:

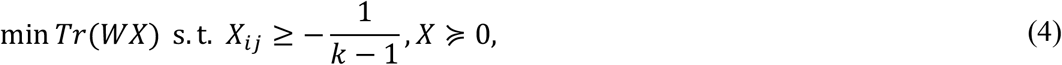

in which *X* ≽ 0 indicates that *X* is a positive semi-definite matrix. Note that 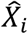 the *i*-th column of the optimum 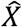, corresponds to the *i*-th fragment. The matrix 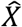 is used to estimate the cluster membership of each fragment using a randomized approach [19]. Each fragment is assigned to the *k*-th cluster when the corresponding column is the closest to the *k*-th random vector in terms of inner product [20]. To do so, firstly, *K* random vectors {*v*_1_, … , *v*_*K*_} are generated, each an *N* × 1 vector with elements drawn from a standard normal distribution. Next, inner products between columns of 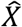 and these random vectors are calculated and the *i*-th fragment is assigned to the *k*-th cluster, corresponding to the *k*-th haplotype, if

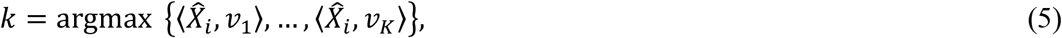

in which 〈.,. 〉 represents the inner product of two vectors.

We exploit dual theory in optimization to solve the semidefinite programming problem (3). Note that the identity matrix is a positive definite matrix, and all its elements are nonnegative. Thus, the identity matrix belongs to the interior of the optimization domain. Thus, the optimization is strictly feasible. Therefore, Slater’s condition is satisfied for the optimization, which immediately results in strong duality (section 5.2.3 of [21]). To derive the dual optimization problem of (4), the Lagrangian function can be written as

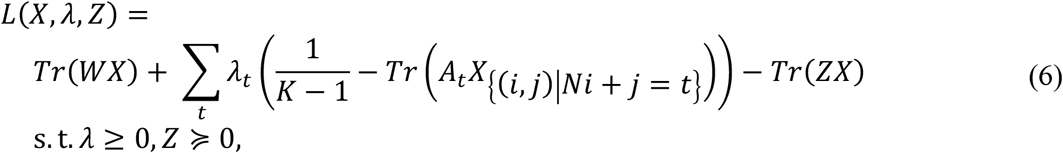

in which *A*_*t*_ is a matrix with the same dimensions as *X* of zeroes with a 1 in the (*i*, *j*)-th element. Then, (6) can be rearranged to

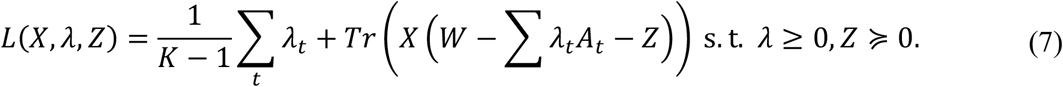

Since the second term is affine in *X*, we should make it bounded. To this end, the weight of the affine function should be zero. Thus, the maximization (6) can be simplified to

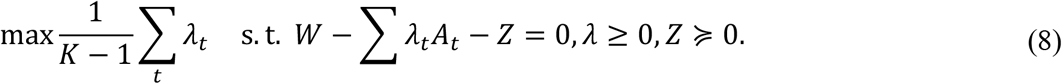

To achieve an unconstrained optimization, we define the augmented Lagrangian function of the optimization as [22]:

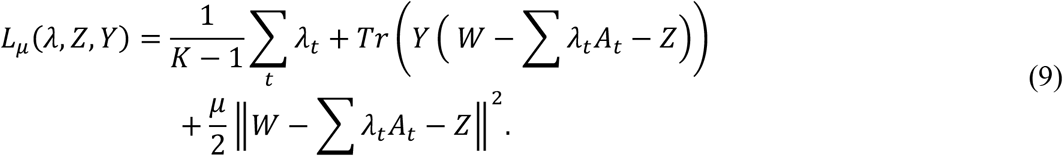

A novel iterative optimization scheme for solving the max-*K*-cut problem then becomes:

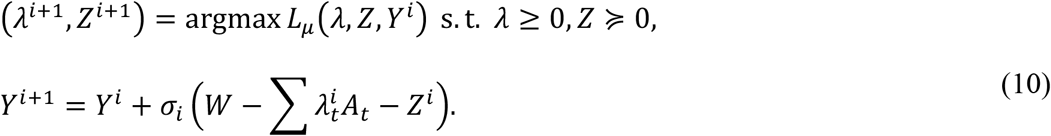

Then, the optimality condition of the first optimization results in a linear equation, which is solved by a Newton conjugate gradient approach (Section 10.2 of [23]). We stop the iteration when the relative duality gap (defined as 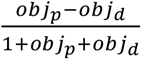 in which *obj*_*p*_ and *obj*_*d*_ are the value of primal and dual objective functions, respectively [26]) falls below a certain convergence threshold, which we set to 0.01. Note that the smaller this threshold, the longer the runtime but the better the estimate (Supplementary information: Table S2). Then, the primal optimal point *X* is found using complementary slackness conditions (section 5.5.2 of [21]). To implement the mentioned algorithm, we use the SDPNAL+ package [26].

### 2.4 Experimental setup

#### 2.4.1 Data

In order to evaluate the performance of the developed pipelines and algorithms, we consider numerous scenarios on both simulated and experimental data. First, we performed extensive simulation experiments using the reference genome of potato (*Solanum tuberosum)* as a basis. We first simulated data based on a region of one million base pairs (1 Mb) starting from position 5,032,020 on chromosome 1 and subsequently used the full chromosome 1 sequence (88.6 Mb). We introduce SNPs in the reference at a rate of one per 100 or 1000 (for the 1MB region) and one per 100 for the full chromosome. We generate synthetic triploid and tetraploid genomes as FASTA files by combining *K* = 3 resp. *K* = 4 mutated copies of the reference sequence using the haplo-generator routine from the Haplosim package [4]. This package also produces *K* true haplotypes in a text file, including the genomic positions of SNPs and the corresponding alleles, which are used for evaluation (see 2.4.2).

Subsequently, we simulated several linked-read datasets following the 10X technical specifications, using the LRSIM package [24]. We set the number of molecules per bead (-m) as 10 and assigned the number of barcodes (-t) such that the molecule coverage is 0.2, as discussed in the 10X Genomics technical note (No. CG00044). The output of each LRSIM simulation consists of two FASTQ files, containing paired-end reads with length of 2×151 bp, in which the first 16 bases are the barcode sequence. The outer distance between the two reads in a pair is set to the default value, 350, with a standard deviation of 35. Then, as described in Section 1.1, the LongRanger and FreeBayes packages are used for aligning reads and calling SNPs, respectively.

To the best of our knowledge, there is no publicly available, real dataset for a polyploid organism containing true haplotype sets, which makes it hard to determine accuracy. To obtain an impression of the distribution of haplotype block lengths and runtimes, we download 10X raw read data of hexaploid sweet potato (*Ipomoea batatas*) from the NCBI database (accession SRX4706082) [25].

#### 2.4.2 Performance assessment

To evaluate the length of the reconstructed haplotypes, we calculate and report the mean value over all haplotype blocks. To assess the accuracy of each algorithm, we consider two criteria: reconstruction rate, a measure of local accuracy; and vector error rate, a more global measure. Given reconstructed haplotypes 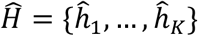 and ground truth haplotypes *H* = {*h*_1_, … , *h*_*k*_}, the reconstruction rate is defined as:

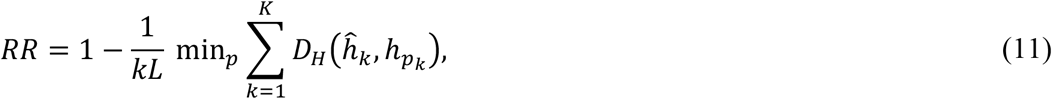

in which *L* is the haplotype length and *D*_*H*_(.,.) is the Hamming distance function, which counts the number of mismatch elements between its arguments. Additionally, *p* is a permutation on the set {1, … , *K*}, and *p*_*k*_ is the *k*-th element of *p*. We calculate this criterion for each haplotype block and report the average. The vector error rate is calculated by finding the minimum number of switches needed in haplotype segments in order to match 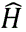 to *H*; this number is then divided by the haplotype length [7, 15].

Since for real data there is no ground truth for assessing the performance of the estimated haplotype, the mentioned metrics cannot be used. To handle this issue, another metric, the Minimum Error Correction (MEC) score, has been frequently used in the literature [5–6]:

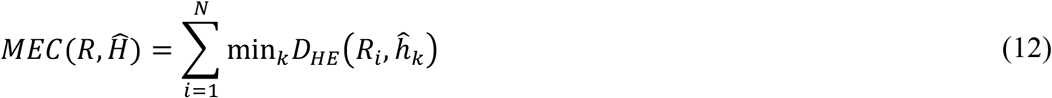

in which *R*_*i*_ is the *i*-th pre-processed read (Section 2.1). For haplotypes with a length of *l*, the extended Hamming distance function is defined as 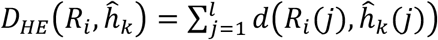. The value 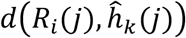 will be one when read *R*_*i*_ covers the *j*-th position of haplotype 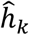 and both are of the same allele, and will be zero otherwise. To interpret this metric, we should note that MEC shows the extent of match between the reconstructed haplotypes and the read dataset.

## 3. Results

We have developed Hap10, a novel pipeline for haplotyping polyploids based on 10X linked-read data. The basis of Hap10 is a set of pre-processing steps called Hap++. After application of Hap++, SDhaP [6] can be used as an assembly core. We also propose an alternative core, based on the SDPNAL+ algorithm. The combination of Hap++ and the new assembly core is called Hap10.

To obtain an impression of the performance of SDhaP, Hap++/SDhaP and Hap10, we performed extensive simulations based on real-world data, the potato genome. This allows us to investigate accuracy (Section 2.4.2) and run time in different scenarios, varying sequence length, coverage, ploidy, heterozygosity etc. (Section 3.1). We then apply the pipeline to real-world data to evaluate performance in terms of haplotype block length and run time (Section 3.2).

### 3.1 Simulated data

We first applied the various algorithms on 10X data simulated based on a relatively short stretch of the potato genome, of 1 Mb (section 2.4.1), to learn about the influence of various genome and sequencing characteristics.

#### 3.1.1. Linked-read information yields longer haplotypes

As a first test, we applied SDhaP to the simulated read data with and without taking the barcode information into account. The program has no problem dealing with data for a region of this length. Without linked-read information, the reconstruction rate and the vector error rate are relatively good, but the reconstructed haplotype blocks are very short, 11.8 SNPs on average (Supplementary information: Table S3, first row) as is to be expected.

Taking the linked read information into account here improves average haplotype block length dramatically, to over 6,000 SNPs (Supplementary information: Table S3, second row compared to the first row). At the same time, the reconstruction rate drops, and the vector error rate increases, indicating low quality haplotypes. This is due to the effect of mixed haplotypic origin of fragments, misleading the haplotype assembly program. It can be also considered the consequence of the poor connections between subgraphs, insufficient for haplotyping, as illustrated in Supplementary information: Figure S1. An approach in which haplotypes are calculated independently on three equally sized parts of the region of interest supports this: the average block length decreases, but both reconstruction rate and vector error rate improve (Supplementary information: Table S3, third row compared to the second row). This suggests that while SDhaP in principle works for haplotype assembly in polyploids, performance may be improved by pre-processing the data.

#### 3.1.2. Preprocessing by Hap++ yields shorter, more reliable haplotype blocks

To solve the problems encountered in Section 3.1.1, we developed a novel preprocessing pipeline Hap++, to extract strongly connected components from the fragment graph. This reduces the potential for erroneous haplotype assembly, at the expense of a reduced haplotype block length. We apply Hap++ to triploid and tetraploid data simulated on the 1 Mb region taken from the potato genome, at various levels of coverage (2, 5 and 10 per haploid) and different SNP rates (0.01 and 0.001). We repeated the simulations 5 times and report average haplotype block lengths, reconstruction rates and vector error rates in Figure 3.

**Figure 3.**
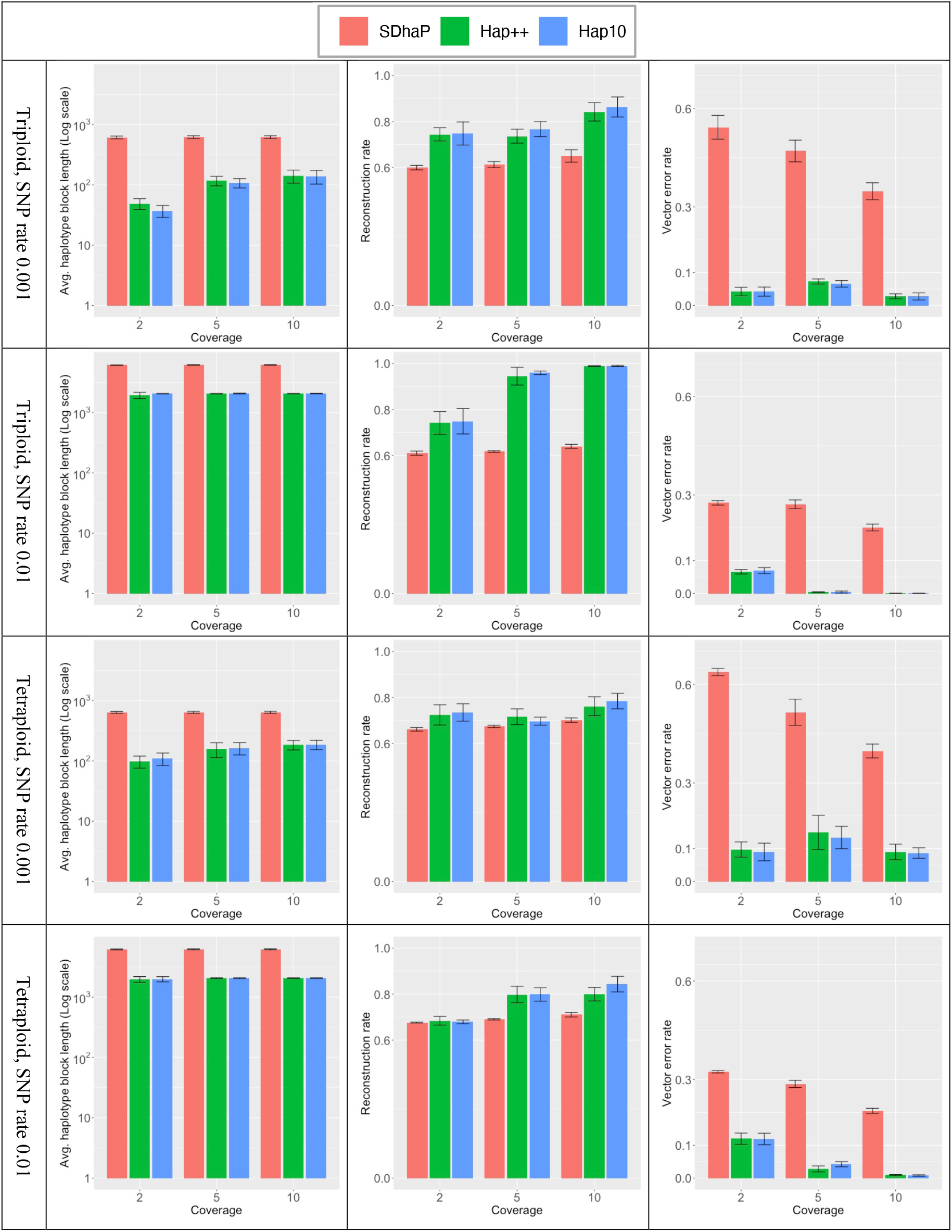
Average haplotype block length (left column, note the logarithmic scale), reconstruction rate (middle) and vector error rate (right) for different coverage levels. Bars indicate averages, whiskers standard deviation of 5 repeated simulations.

Hap++ indeed yields much shorter haplotype blocks (e.g. 339.9 versus 787.2 Kb for SDhaP for triploid, SNP rate 0.01, coverage 10), but drastically improves performance over SDhaP. The reconstruction rate increases, in particular for the triploid simulations, and the vector error rate drops to below 0.1 for almost all simulations where for SDhaP it can reach as high as 0.6. This indicates that the spurious connection problem discussed before occurs in practice and seriously impacts results. It is clear that the SNP rate has a large influence on performance: at low SNP rates, average haplotype block lengths are shorter and accuracy is higher, again particularly for the triploid simulations.

Figure 3 also shows that performance improves with coverage (as expected), and that a coverage of 2 is so low that all methods make errors, due to the fact that SNPs often cannot even be detected. Hap++ benefits more quickly from increasing coverage than SDhaP. SDhaP performance improves up to a coverage of 10 per haploid and keeps improving, as spurious connections in the fragment graph will increasingly be supported by more connections and errors will be counteracted by solid data: in fact, SDhaP needs 5 times as much coverage to reach a similar vector error rate (Supplementary information: Table S4).

Figure 4 shows performance at different ploidy levels. While haplotype block length is invariant to the ploidy level, in most cases more trustworthy haplotypes are attained at higher ploidy levels. To understand this, note that the max-*K*-cut randomized approach (part of the assembly core) is theoretically guaranteed to converge to near the optimal value (by a factor of 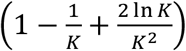, a function increasing in *K*, as presented in Theorem 1 of [19]). However, limited precision in the SDP solver means this solution is not always found in practice.

**Figure 4.**
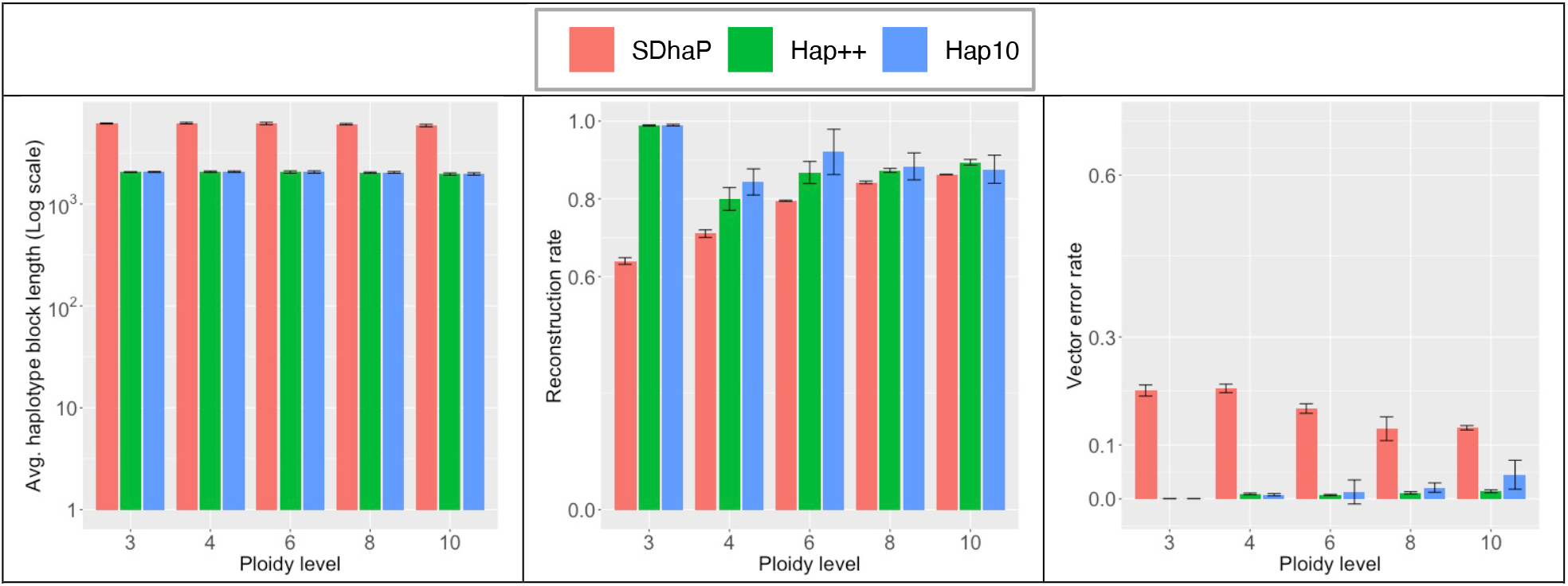
Average haplotype block length (left), reconstruction rate (middle) and vector error rate (right) for different ploidy levels (SNP rate 0.01). Bars indicate averages, whiskers standard deviation of 5 repeated simulations.

#### 3.1.3. Hap++ deals better with imperfect 10X data

Ideally, the 10X technology ensures each unique barcode is assigned to fragments that originate from a single, long DNA molecule. In practice however, fragmentation is imperfect, leading to shorter molecules, and more than one molecule may receive the same barcode (see Section 2.2). Hap++ contains a pre-processing step to cluster reads based on the expected molecule size, to avoid the concatenation of different molecules in a single line of the fragment file as much as possible.

Figure 5 (top) shows performance as a function of both the number of molecules that on average receives the same barcode (in simulated data). The difference between SDhaP and Hap++ is striking, in that vector error rate increases drastically with the number of molecules per barcode for SDhaP but remains negligible for Hap++. The Hap++ reconstruction rate decreases somewhat, but remains higher than that of SDhaP up to at least 10 molecules per barcode – which, given the sequence length of 1 Mb and the molecule length of 100 kb entails a significant probability of overlap between molecules with the same barcode.

**Figure 5.**
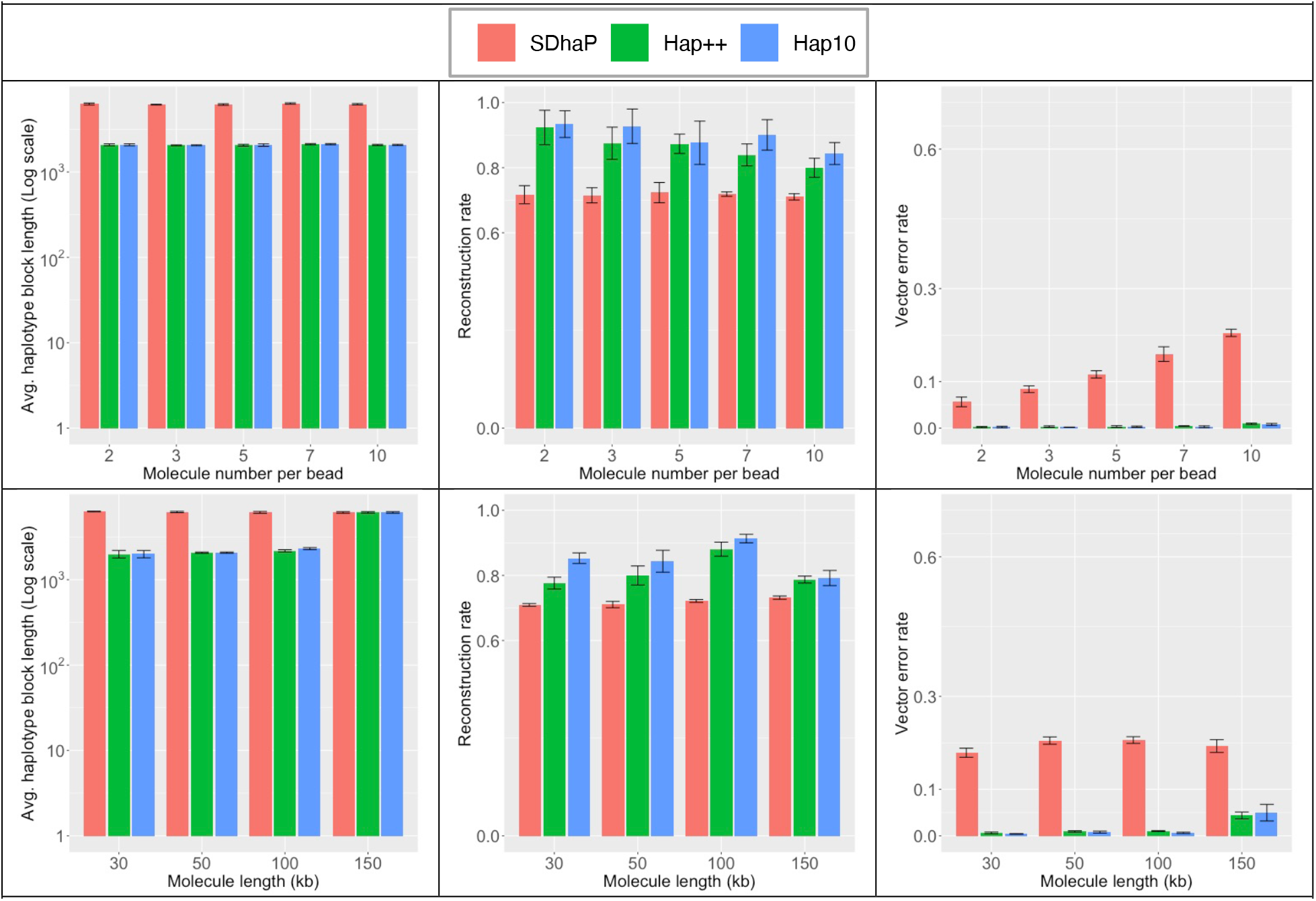
Average haplotype block length (left), reconstruction rate (middle) and vector error rate (right) for different settings of the 10X linked-read simulation (SNP rate 0.01, tetraploid), varying the number of molecules per bead (top) and the molecule length (bottom). Bars indicate averages, whiskers standard deviation of 5 repeated simulations.

We also varied the length of the 10X molecules in the simulations, from 30, 50 and 100 to 150 kb. Figure 5 (bottom) shows that longer molecules yield better haplotypes in terms of reconstruction rate due to the improved long-range information, but eventually increases the vector error rate, likely due to the increased probability of overlap of such long molecules (150 kb in a 1 Mb region).

#### 3.1.4. Hap10 improves performance, at considerable computational cost

Figure 3 also includes performance of Hap10, a combination of the Hap++ pre-processing stage with a new assembly core based on the SDPNAL+ algorithm. Overall, Hap10 and Hap++ perform more or less on par, with a slight advantage for Hap10 at higher coverage levels, at lower molecule lengths and when more molecules receive the same barcode. This suggests the Hap10 assembly core is more robust to errors and problems due to imperfect 10X data. However, this comes at a cost: the Hap10 runtime is significantly higher. Table 1 reports CPU times for the results reported in Figure 3. The pipelines were run on 24 CPU cores of a machine with 48 cores (Intel Xeon Silver 4116) and 754 GiB system memory. Clearly, the pre-processing by Hap++ occurs a time penalty, most visible for lower coverages, which pays off in a quicker runtime of the final SDhaP application, clearly seen at higher coverages. Hap10 is up to two orders of magnitude slower. When this is worth the effort, the pipeline can be run in *accurate mode* (using Hap10 optimization) with high haplotype quality, or in *fast mode* (using Hap++) with reasonable quality, depending on user preference.

**Table 1.** Run times (seconds) of the algorithms compared in Figure 3.

| Ploidy | SNP rate | Coverage | SDhaP | Hap++ | Hap10 |
| --- | --- | --- | --- | --- | --- |
| Triploid | 0.001 | 2 | 4 | 53 | 419 |
|  |  | 5 | 24 | 54 | 2,919 |
|  |  | 10 | 34 | 92 | 1,590 |
|  | 0.01 | 2 | 3 | 64 | 2,590 |
|  |  | 5 | 150 | 162 | 8,590 |
|  |  | 10 | 660 | 400 | 19,221 |
| Tetraploid | 0.001 | 2 | 8 | 39 | 710 |
|  |  | 5 | 21 | 90 | 4,119 |
|  |  | 10 | 65 | 204 | 13,824 |
|  | 0.01 | 2 | 43 | 98 | 15,307 |
|  |  | 5 | 478 | 370 | 28,598 |
|  |  | 10 | 1,497 | 1,022 | 36,736 |

#### 3.1.5. Hap++ and Hap10 work on longer sequences

As a final test, we generated linked read data for the full chromosome 1 of the potato genome, simulating a tetraploid genome at a SNP rate of 0.01. The coverage is 10 per haploid genome. Results are reported in Table 2. Notably, SDhaP encountered a segmentation fault in this simulation, leaving us unable to report a result. Hap++ and Hap10 provide haplotypes with the same block lengths, with better accuracy in terms reconstruction rate and vector error rate. Moreover, the MEC between the read set and the reconstructed haplotypes is lower, suggesting a better compatibility between the two. However, as before, the computational cost of Hap10 is significant at approx. 900 CPU hours vs. 12 hours for Hap++.

**Table 2.** Results for chromosome 1 of a tetraploid potato with coverage 10 per haploid and a SNP rate of 0.01.

| Method | Avg.<br>haplotype<br>block length<br>(no. SNPs) | N50<br>haplotype<br>block length<br>(bp) | Reconstru<br>ction rate | Vector<br>error rate | MEC | CPU<br>time<br>(min) |
| --- | --- | --- | --- | --- | --- | --- |
| SDhaP | - | - | - | - | - | - |
| Hap++ | 3,923 | 828,058 | 0.88 | 0.0083 | 342,956 | 741 |
| Hap10 | 3,923 | 828,058 | 0.92 | 0.0070 | 218,635 | 54,835 |

### 3.2 Real data

To obtain an idea of the applicability of Hap++ and Hap10 to real data, we ran the pipeline to reconstruct the six haplotypes of chromosome one of sweet potato (with the length of 36 Mb) based on 10X data available in the NCBI Short Read Archive.

The length distribution of the reconstructed haplotypes is displayed in Figure 6; the N50 length of the blocks is 78.4 resp. 78.3 kb for Hap++ and Hap10. The MEC scores between the read set and the reconstructed haplotypes are 122,363 resp. 133,282 for the reconstructed haplotypes using Hap++ and Hap10, respectively, which would indicate that in this case the reconstructed haplotypes by Hap++ are more compatible with the read dataset than those generated by Hap10. However, true accuracy can only be evaluated by comparison to a ground truth.

**Figure 6.**
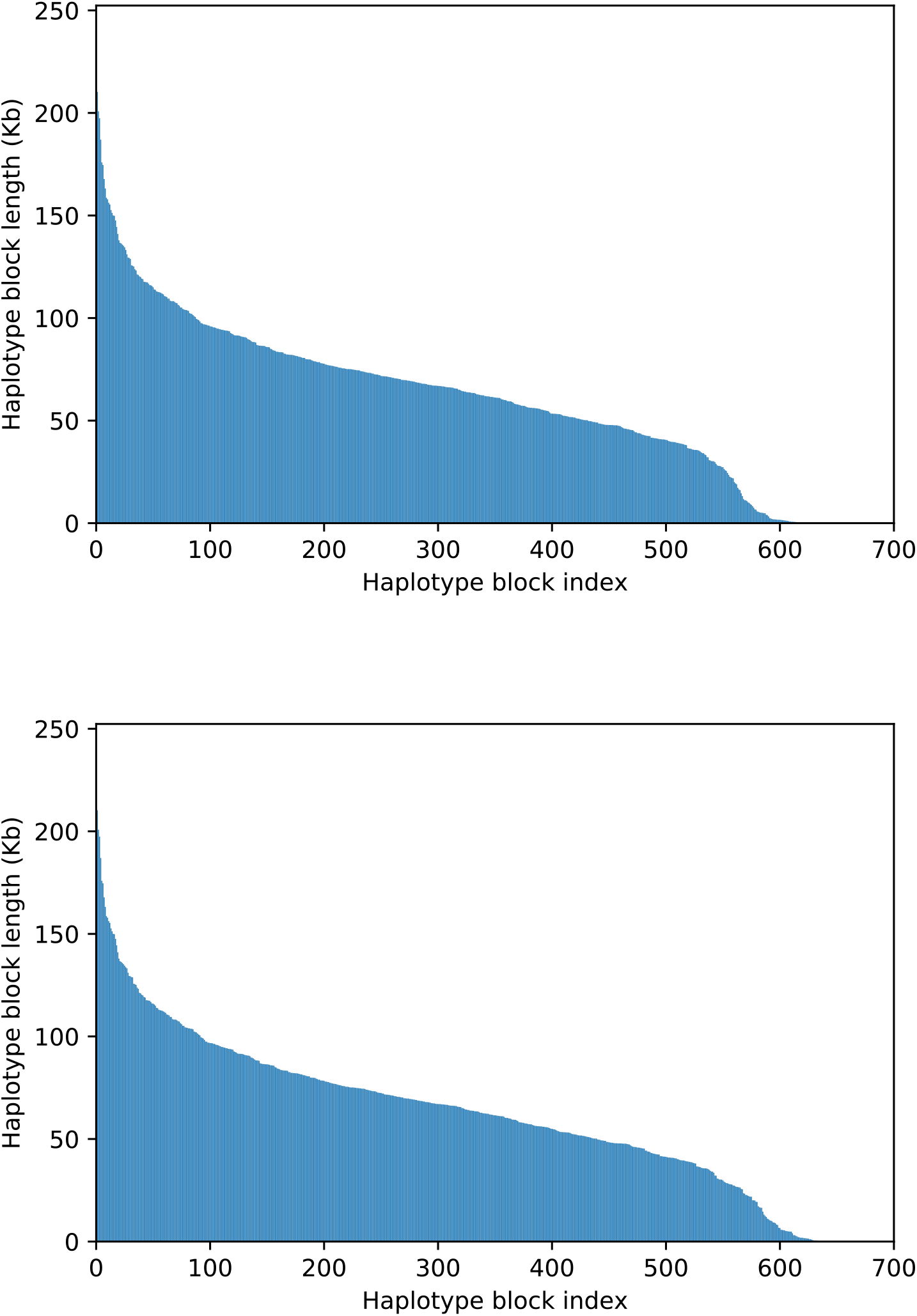
Haplotype block length distributions for 10X real data of sweet potato using Hap++ (top) and Hap10 (bottom).

## 4. Conclusion

We developed a first haplotyping pipeline specifically for 10X linked-read data generated from a polyploid genome. It makes haplotyping full chromosomes of complex genomes feasible. The proposed Hap++ preprocessing pipeline improves on the accuracy of immediate application of SDhaP by approximately 30% (resp. 20%) on simulated 10X data of triploids (resp. tetraploids) at the cost of a decreased haplotype block length. Our framework builds on SDhaP, a typical Illumina haplotyping algorithm, using a standard fragment file as input. Any improvement in SDhaP or similar algorithms thus may immediately enhance 10X haplotyping. The proposed novel optimization scheme, Hap10, provides even more accurate haplotypes, albeit at significant computational cost.

One topic for future research is to consider different optimization techniques for the max-*K*-cut clustering problem [27]. A new method based on linear programming [28] may provide a solution for overcoming the high runtime involved in the semidefinite programming problem. A second avenue for research is automatic optimization of the key parameters of the pipeline, specifically the threshold *t* for the normalized cut algorithm and the convergence threshold used in the optimization step.

## Declarations

### Ethics approval and consent to participate

Not applicable.

### Consent for publication

Not applicable.

### Availability of data and materials

- Reference genome of *Solanum tuberosum*: ftp://ftp.ensemblgenomes.org/pub/plants/release-42/fasta/solanum_tuberosum/dna/
- 10X read data of sweet potato: https://www.ncbi.nlm.nih.gov/sra/SRX4706082
- LRSIM: https://github.com/aquaskyline/LRSIM
- LongRanger: https://github.com/10XGenomics/longranger
- FreeBayes: https://github.com/ekg/freebayes
- SDPNAL+: https://blog.nus.edu.sg/mattohkc/softwares/sdpnalplus/
- Scikit-learn: https://scikit-learn.org/
- SDhaP: https://sourceforge.net/projects/SDhaP/
- Our pipeline and code: https://github.com/smajidian/Hap10

### Authors’ contributions

Methods and experiments were designed by SM and DdR. Algorithm code was implemented by SM. SM and DdR wrote the manuscript. DdR and MHK supervised the project. All authors read and approved the final manuscript.

## Acknowledgements

We would like to thank Brian Lavrijssen for helpful discussions.

## Supplementary information

**Figure S1.**
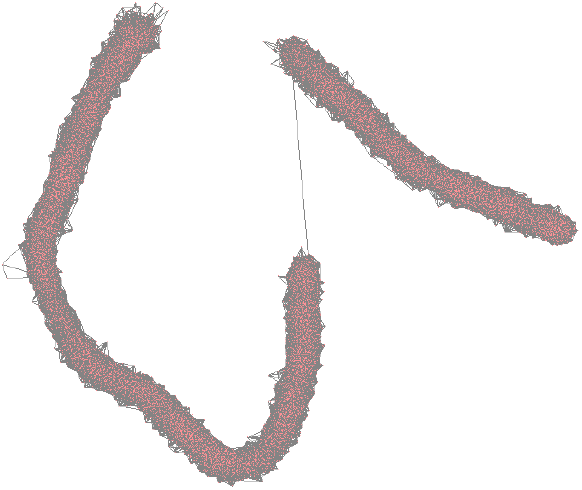
A graph indicating overlap between fragments. Red dots are vertices (corresponding to the fragments), grey lines are edges drawn when two fragments have at least one SNP in common. The depicted graph is for a case with 5 mb reference genome containing an N-region of 50kb. The coverage is 15 per haploid and the SNP rate is 0.01. The average length of 10X DNA molecules for this simulation is set to 50kb. Few fragments originate from a DNA molecule larger than 50kb. The resulting graph has two separate subgraphs connected by a single edge. Note that one barcode-specific fragment connecting two read blocks is not sufficient for connecting the corresponding haplotypes. This phenomenon decreases the quality of reconstructed haplotype. The figure is generated using Cytoscape (www.cytoscape.org).

**Table S1.** An example of the haplotype output format. We report the reconstructed haplotypes as a text file with a specific format similar to that of HapCUT2. Each haplotype block starts with a line describing the length of the haplotype, number of reads corresponding to the block and the minimum error correction (MEC) score. From the next line, each row corresponds to each variant. The first and second columns show the 1-based index and variant position, respectively. Then, the next 2 * ploidy columns are haplotypes and quality scores. For each allele of haplotypes, a quality score is provided. As a metric for quality, we use the number of matching reads at each position that are estimated for each haplotype.

| Block 1 | Length of haplotype block 9 |  |  |  | Number of read 58 |  | Total MEC 10 |
| --- | --- | --- | --- | --- | --- | --- | --- |
| 1 | 644 | 0 | 0 | 1 | 2 | 9 | 8 |
| 2 | 783 | 0 | 0 | 1 | 10 | 13 | 15 |
| 3 | 916 | 0 | 0 | 1 | 9 | 16 | 13 |
| 4 | 990 | 0 | 0 | 1 | 11 | 12 | 16 |
| 5 | 1848 | 0 | 1 | 0 | 13 | 13 | 12 |
| 6 | 1862 | 1 | 0 | 0 | 13 | 13 | 14 |
| 7 | 1930 | 1 | 0 | 1 | 11 | 11 | 12 |
| 8 | 1042 | 0 | 0 | 1 | 13 | 14 | 16 |
| 9 | 1075 | 0 | 0 | 1 | 10 | 16 | 14 |

**Table S2.**
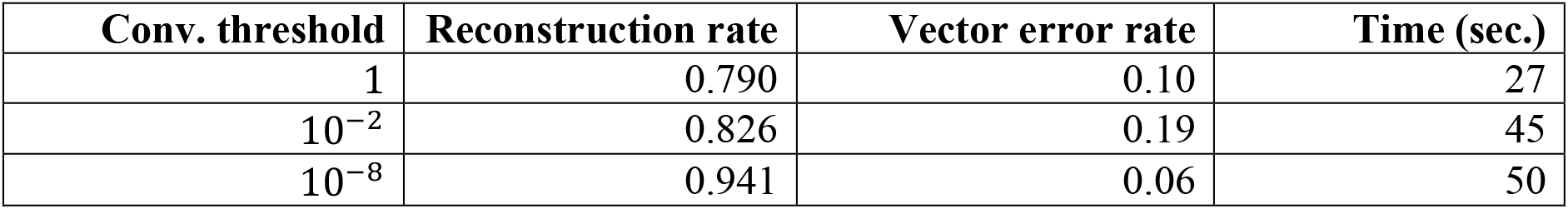
The impact of the convergence threshold on Hap10 performance. A triploid genome of 230kb with a SNP rate of 0.001 is simulated. The average molecule length and number of molecules per bead are 50k and 10, respectively.

**Table S3.**
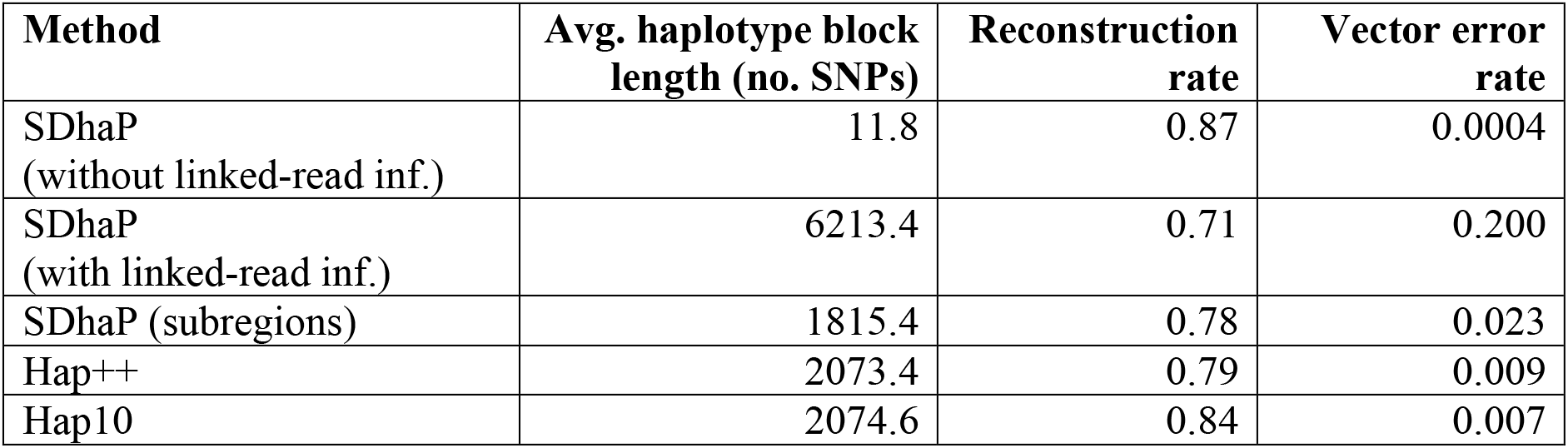
SDhaP with and without linked-read information. For the latter, the input data is considered as regular Illumina reads and barcodes are not used. The dataset is simulated using 1 Mb of chromosome one of potato genome with a SNP rate of 0.01. The coverage is 10. The results are averaged over 5 independent simulations. For the third row, we split the 1 Mb region into three independent parts of the same size. The last two rows present results of Hap++ and Hap10 on the same data.

| **Method** | **Avg. haplotype block length (no. SNPs)** | **Reconstruction rate** | **Vector error rate** |
| --- | --- | --- | --- |
| SDhaP<br>(without linked-read inf.) | 11.8 | 0.87 | 0.0004 |
| SDhaP<br>(with linked-read inf.) | 6213.4 | 0.71 | 0.200 |
| SDhaP (subregions) | 1815.4 | 0.78 | 0.023 |
| Hap++ | 2073.4 | 0.79 | 0.009 |
| Hap10 | 2074.6 | 0.84 | 0.007 |

**Table S4.**
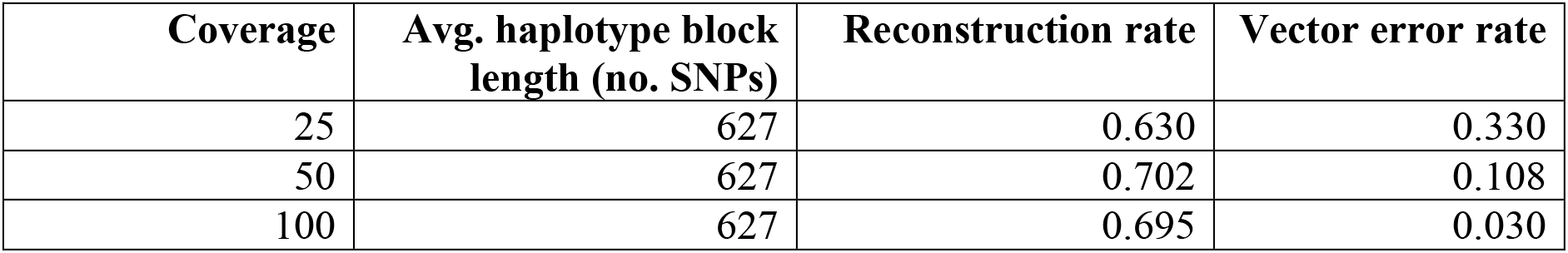
Performance of SDhaP at different coverage levels, for a triploid genome with SNP rate of 0.001. The average molecule length and number of molecules per bead are 50k and 10, respectively. The results are averaged over 5 independent simulations.

| **Coverage** | **Avg. haplotype block length (no. SNPs)** | **Reconstruction rate** | **Vector error rate** |
| --- | --- | --- | --- |
| 25 | 627 | 0.630 | 0.330 |
| 50 | 627 | 0.702 | 0.108 |
| 100 | 627 | 0.695 | 0.030 |

## References

1. Comai L. The advantages and disadvantages of being polyploid. Nature Reviews Genetics. 2005;6(11):836–46.

2. Qian L, Hickey LT, Stahl A, Werner CR, Hayes B, Snowdon RJ, Voss-Fels KP. Exploring and harnessing haplotype diversity to improve yield stability in crops. Frontiers in Plant Science. 2017;8:1534.

3. Liu PY, Zhang YY, Lu Y, Long JR, Shen H, Zhao LJ, et al. A survey of haplotype variants at several disease candidate genes: the importance of rare variants for complex diseases. Journal of Medical Genetics. 2005;42(3):221–7.

4. Motazedi E, Finkers R, Maliepaard C, de Ridder D. Exploiting next-generation sequencing to solve the haplotyping puzzle in polyploids: a simulation study. Briefings in Bioinformatics. 2017;19(3):387–403.

5. Berger E, Yorukoglu D, Peng J, Berger B. HapTree: A novel Bayesian framework for single individual polyplotyping using NGS data. PLoS Computational Biology. 2014;10(3):e1003502.

6. Das S, and Vikalo H. SDhaP: haplotype assembly for diploids and polyploids via semi-definite programming. BMC Genomics. 2015;16:260.

7. Xie M, Wu Q, Wang J, Jiang T. H-PoP and H-PoPG: heuristic partitioning algorithms for single individual haplotyping of polyploids. Bioinformatics. 2016;32(24):3735–44.

8. Siragusa E, Haiminen N, Finkers R, Visser R, Parida L. Haplotype assembly of autotetraploid potato using integer linear programming. Bioinformatics. 2019;35(21):4534.

9. Goodwin S, McPherson JD, McCombie WR. Coming of age: ten years of next-generation sequencing technologies. Nature Reviews Genetics. 2016;17(6):333–51.

10. Weisenfeld NI, Kumar V, Shah P, Church DM, Jaffe DB. Direct determination of diploid genome sequences. Genome Research. 2017;27(5):757–67.

11. Marks P, Garcia S, Barrio AM, Belhocine K, Bernate J, Bharadwaj R, et al. Resolving the full spectrum of human genome variation using linked-reads. Genome Research. 2019;29(4):635–45.

12. Edge P, Bafna V, Bansal V. HapCUT2: robust and accurate haplotype assembly for diverse sequencing technologies. Genome Research. 2017;27(5):801–12.

13. Majidian S, Kahaei MH. NGS based haplotype assembly using matrix completion. PLoS ONE, 2019;14(3):e0214455.

14. Garrison E, Marth G. Haplotype-based variant detection from short-read sequencing. arXiv:1207.3907. 2012.

15. Motazedi E, de Ridder D, Finkers R, Baldwin S, Thomson S, Monaghan K, Maliepaard C. TriPoly: haplotype estimation for polyploids using sequencing data of related individuals. Bioinformatics. 2018;34(22):3864–72.

16. Comaniciu D, Meer P. Mean shift: a robust approach toward feature space analysis. IEEE Transactions on Pattern Analysis & Machine Intelligence. 2002;24(5):603–19.

17. Pedregosa F, Varoquaux G, Gramfort A, Michel V, Thirion B, Grisel O, et al. Scikit-learn: Machine learning in Python. Journal of Machine Learning Research. 2011;12:2825–30.

18. Shi J, Malik J. Normalized cuts and image segmentation. IEEE Transaction on Pattern Analysis and Machine Intelligence. 2000;22(8):888–905.

19. Frieze A, Jerrum M. Improved approximation algorithms for max *k*-cut and max bisection. Algorithmica. 1997;18(1):67–81.

20. de Klerk E, Pasechnik DV, Warners JP. On approximate graph colouring and max-*k*-cut algorithms based on the θ-function. Journal of Combinatorial Optimization. 2004;8(3):267–94.

21. Boyd S, Vandenberghe L. Convex optimization. Cambridge University Press; 2004.

22. Rockafellar RT. Augmented Lagrangians and applications of the proximal point algorithm in convex programming. Mathematics of Operations Research. 1976;1(2):97–116.

23. Golub GH, Van Loan CF. Matrix computations. Johns Hopkins University Press; 1996.

24. Luo R, Sedlazeck FJ, Darby CA, Kelly SM, Schatz MC. LRSim: a linked reads simulator generating insights for better genome partitioning. Computational and Structural Biotechnology Journal. 2017;15:478–84.

25. Wu S, Lau KH, Cao Q, Hamilton JP, Sun H, Zhou C, et al. Genome sequences of two diploid wild relatives of cultivated sweetpotato reveal targets for genetic improvement. Nature Communications. 2018;9(1):4580.

26. Yang L, Sun D, Toh KC. SDPNAL++: a majorized semismooth Newton-CG augmented Lagrangian method for semidefinite programming with nonnegative constraints. Mathematical Programming Computation. 2015;7(3):331–66.

27. Ghaddar B, Anjos MF, Liers F. A branch-and-cut algorithm based on semidefinite programming for the minimum *k*-partition problem. Annals of Operations Research. 2011;188(1):155–74.

28. de Sousa VJR, Anjos MF, Le Digabel S. Improving the linear relaxation of maximum *k*-cut with semidefinite-based constraints. EURO Journal on Computational Optimization. 2019;7(2):123–51.

